# PhoPQ regulates quorum sensing in small colony variant subpopulations of *Pseudomonas aeruginosa*

**DOI:** 10.64898/2026.05.10.723372

**Authors:** Kayla A. Simanek, Alicia G. Mendoza, Amanda F. Kurtz, Autumn N. Pope, Megan L. Schumacher, Jon E. Paczkowski

## Abstract

*Pseudomonas aeruginosa* is an opportunistic human pathogen that is hospital-endemic, forming biofilms on medical equipment and causing thousands of hospital-acquired infections each year. The success of *P. aeruginosa* as an opportunistic pathogen is linked to its phenotypic and genotypic adaptability. *P. aeruginosa* coordinates population-level decisions using quorum sensing (QS), a mechanism of cell-cell communication, and forms small colony variant (SCV) subpopulations in response to stresses such as oxygen limitation and antibiotic exposure. Here, we show that QS mutations promote phenotypic variation; specifically, deletion of *lasR* and autoinducer-modulating mutations in *rhlI* enhance SCV formation in a time-dependent manner, and the resulting SCV subpopulations exhibit distinct biofilm traits. Using transcriptome analysis of isogenic SCV and large colony variant (LCV) populations, we identify the PhoPQ two-component signal transduction system as upregulated in SCVs and show that PhoPQ governs SCV formation in a magnesium-dependent manner. Our data further demonstrate that unphosphorylated PhoP represses *rhlR* gene expression, and that PhoQ-dependent phosphorylation of PhoP de-represses QS to drive virulence factor production and the SCV phenotype in a *lasR* background. Together, these findings provide insight into how clinically evolved QS mutations may serve as a bet-hedging strategy to promote SCV formation and alter QS signaling within a subpopulation.

## INTRODUCTION

Several bacterial human pathogens undergo phase variation and form physiologically distinct subpopulations with different virulence characteristics [1–6]. *P. aeruginosa* forms a small colony variant (SCV) phenotype under stress conditions that results in hyper-biofilm formation due to increased production of exopolysaccharides. Environmental, clinical, and laboratory strains of *P. aeruginosa* have all been shown to produce SCVs [7, 8], indicating that this phenotype is not unique to a specific niche. Stable chromosomal inversions in the 16S rRNA gene [9] and mutations in cyclic-di-GMP synthesis pathways [10] have been identified as genetic drivers of the SCV phenotype, while antibiotic exposure and oxygen limitation act as selective pressures for SCV formation [11].

In addition to environmental stresses that can induce phenotypic variation, clinical strains of *P. aeruginosa* are well-known to evolve mutations in QS pathways that alter virulence gene regulation and promote population heterogeneity [12–15], highlighting the complexity with which *P. aeruginosa* adapts to its environment [16]. QS is a mechanism of cell-cell communication that relies on the production and detection of small molecules called autoinducers (AIs) [17]. The *P. aeruginosa* QS network is best described as a web of interconnected pathways comprising the Las, Rhl, and Pqs systems [18]. The Las (LasI/R) and Rhl (RhlI/R) systems belong to the well-characterized LuxI/R family [19], in which a synthase (LuxI homolog) produces an acyl homoserine lactone (AHL) AI [20–23] that binds to its cognate transcription factor receptor (LuxR-type homolog) to regulate gene expression [24–33]. The Pqs system is positively regulated by Las and negatively regulated by Rhl to produce quinolone signals for downstream signaling [34–39]. Mutations in *lasR* occur at high frequency in clinical isolates [12, 13, 40–44]. LasR sits atop the canonical QS hierarchy and was initially thought to be primarily responsible for upregulating downstream QS systems, including the Rhl pathway. However, clinical *lasR* mutations often produce a non-functional LasR protein, yet QS proceeds through RhlI/R signaling [12, 14]. RhlI is the AI synthase that produces *N*-butanoyl-L-homoserine lactone (C4HSL), the cognate ligand for the transcription factor RhlR [29, 32]. We showed previously that clinically evolved mutations in *rhlI*; specifically, the amino acid substitutions G62S, D83E, and P159S can alter the timing of virulence phenotypes in a *lasR* strain through re-calibration of AI concentrations in the population [13].

QS networks also integrate signals from other environmental sensing systems, creating crosstalk between diverse regulatory inputs. For example, PhoB/R is a two-component signal transduction system (TCS) that primarily regulates phosphate transport [45, 46]. PhoR is a periplasmic kinase that phosphorylates the cytoplasmic transcription factor PhoB; phosphorylated PhoB upregulates *rhlR* expression [47–49], leading to downstream activation of QS-regulated traits. Additional TCS systems co-regulate QS-dependent traits such as biofilm morphology and alginate production [50]. For example, RhlR directly upregulates *algB*, which encodes a response regulator phosphorylated by BphP in response to light, leading to repression of biofilm formation; dephosphorylation of AlgB by KinB inactivates this repression [51]. Together, these examples illustrate how the gene targets of multiple TCS systems in *P. aeruginosa* overlap with the QS regulon to integrate diverse sensory cues.

Here, we show that *lasR* mutations promote the formation of a stable SCV subpopulation in *P. aeruginosa* liquid cultures, and that autoinducer-modulating mutations in *rhlI* accelerate and enhance this phenotype. The resulting SCV subpopulations exhibit robust biofilm formation and tolerance to polymyxin B. Transcriptomic comparison of SCV, and their large colony variant (LCV) counterpart reveals upregulation of the *phoPQ* genes and downregulation of *rhlIR* in SCVs. The PhoPQ TCS responds to environmental Mg^2+^ levels in *P. aeruginosa* [52–54]; under low magnesium conditions, the transmembrane histidine kinase PhoQ controls the phosphorylation state of the transcription factor PhoP, which upregulates genes involved in maintaining membrane integrity [52, 53, 55, 56]. We show that PhoQ-dependent phosphorylation of PhoP de-represses the *rhlR* promoter, and that the PhoPQ system governs both SCV formation and RhlR-dependent virulence factor production in a *lasR* deletion background. The PhoPQ system has also been linked to polymyxin antibiotic resistance and interactions with epithelial cells [57–59], underscoring its importance in pathogenesis. This study therefore reveals a previously unrecognized regulatory relationship between the PhoPQ TCS and the *P. aeruginosa* QS system.

## RESULTS

### Mutations in *lasR* and *rhlI* promote SCV formation and associated phenotypes

We previously identified clinically evolved *rhlI* mutations in acute strains of *P. aeruginosa* that differed from the laboratory strain UCBPP-PA14. These RhlI variants resulted in re-calibration of AI levels and a restoration of virulence phenotypes in a Δ*lasR* background, specifically [13]. We showed that a Δ*lasR* strain exhibited early and elevated production of C4HSL, which we hypothesize is through metabolic flux driving all AHL substrates into the production of C4HSL instead of them being shared with 3OC12HSL. The early increase in C4HSL levels resulted in a significant lag in RhlR-dependent pyocyanin production, a phenazine known for its redox sensitive properties, which acts as a key virulence factor across host systems. The RhlI variants caused a reduction in C4HSL concentrations to restore timely RhlR activation and downstream upregulation of phenazine biosynthesis genes. We expanded our search for new RhlI variants to include chronic clinical isolates from cystic fibrosis patients and identified a point mutation in *rhlI* encoding the amino acid substitution R41K. We introduced this mutation chromosomally in UCBPP-PA14 in a Δ*lasR* background and assayed for pyocyanin production in defined minimal media (FDS). It was previously shown that FDS media can drive QS-regulated virulence phenotypes in Δ*lasR* strains sooner than rich media. The Δ*lasR* strain expressing the R41K variant produced pyocyanin near wild-type (WT) levels within 5 hours of initiating the time course. At 18 hours pyocyanin production in a Δ*lasR* RhlI R41K background exceeded that of WT, while a Δ*lasR* strain did not produce detectable levels of pyocyanin (**Figure 1A**).

**Figure 1.**
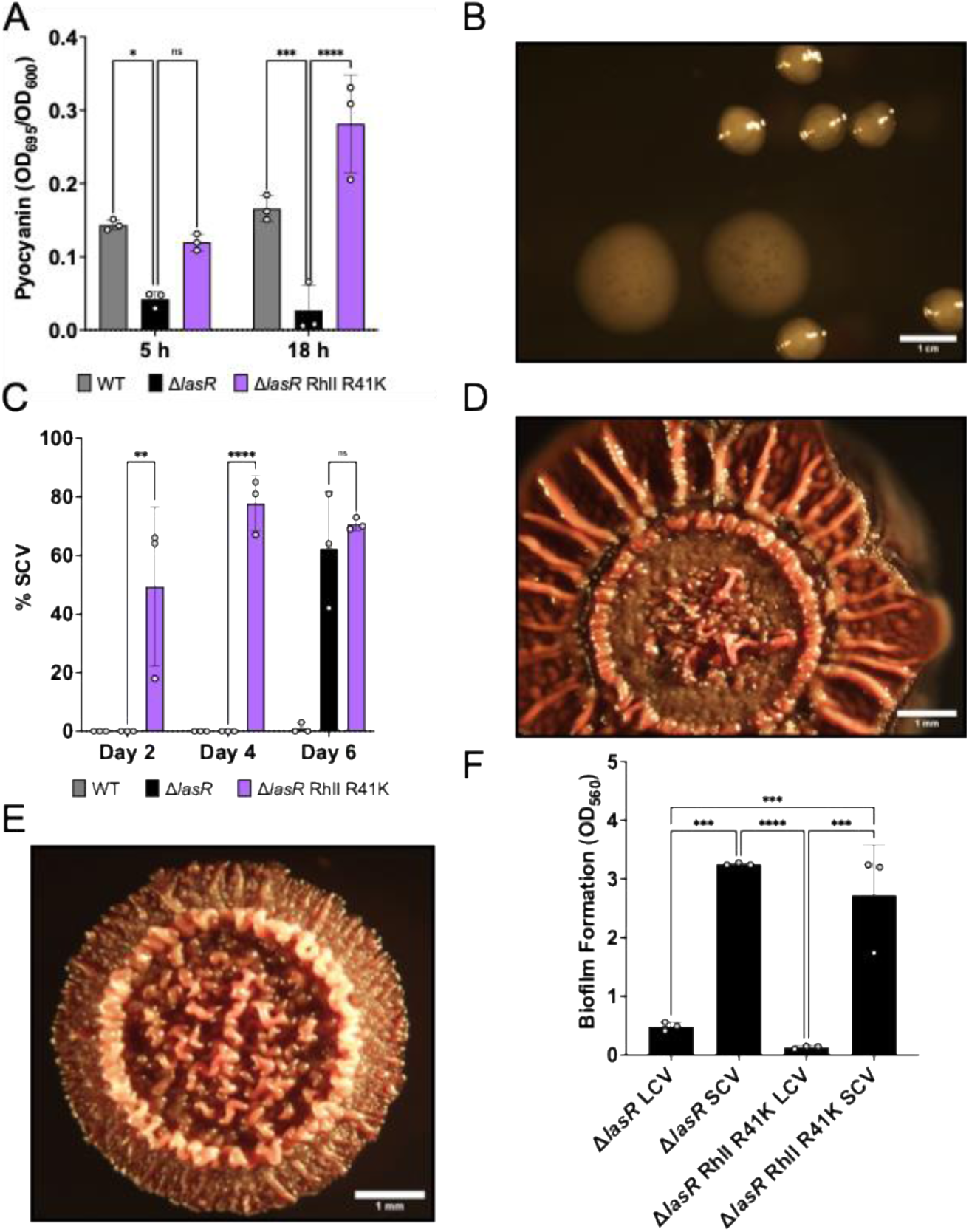
Quorum sensing mutations promote SCV formation and associated phenotypes. (**A**) Pyocyanin production of WT, Δ*lasR*, and Δ*lasR* RhlI R41K strains in FDS minimal media over 18 hours. (**B**) Representative microscopy images of SCVs (arrow) and LCVs. (**C**) SCV production of strains passaged in LB for 6 days. (**D**) Representative image of an LCV colony biofilm on Congo red agar. (**E**) Representative image of a Δ*lasR* RhlI R41K SCV colony biofilm on Congo red agar. (**F**) Crystal violet assays were performed to quantify the biofilm formation of SCVs and LCVs. Error bars represent standard deviations of biological replicates. Statistical analyses were performed using a two-way ANOVA with a Tukey’s multiple comparisons tests. P values: ns ≥0.05; * <0.05; ** <0.01; *** <0.001; **** <0.0001.

During the initial propagation of these strains, we observed that Δ*lasR* strains formed a SCV subpopulation when serially grown in LB liquid culture under aeration (**Figure 1B**). This is distinct from prior observations of *P. aeruginosa* SCV formation, which correlated with static growth and pellicle formation [11]. To quantify SCV formation longitudinally, we passaged WT, Δ*lasR*, and Δ*lasR* RhlI R41K strains in LB for one week with daily colony counting and media replacement. WT failed to consistently produce SCVs throughout the time course, while the Δ*lasR* parent strain produced significant levels of SCVs by day 6 (**Figure 1C**). SCV formation was markedly accelerated in the Δ*lasR* RhlI R41K strain, with SCVs detected as early as day 2 (**Figure 1C**). Increased SCV production was also observed in Δ*rhlI* and Δ*rhlI* Δ*lasR* backgrounds, indicating that the loss of *rhl* system function in the absence of *lasR* supports the SCV phenotype (**Supplementary Figure 1A**).

To assess the stability of the SCV phenotype, individual SCVs were picked and re-passaged in LB for one week; reversion was observed only in the Δ*lasR* background at an average rate of less than 20%, indicating that SCV subpopulations derived from these genetic backgrounds are relatively stable after isolation (**Supplementary Figure 1B**). Whole genome sequencing was then performed on SCVs, LCVs, and SCVs that reverted to LCVs to identify unique genetic features. Aligning with previous literature, we noted mutations in *wspA* (PA14_16430), a chemotaxis gene involved in c-di-GMP signaling that promotes biofilm formation [60], consistent with the established role of this locus in *P. aeruginosa* SCV formation [7,11] (**Supplementary Table 1**). Specifically, we observed that Δ*lasR* RhlI R41K SCVs, as well as all revertants, had a missense mutation at *wspA* G1387A (V464I), suggesting this mutation likely drove initial SCV formation. Similarly, all revertants had a secondary deletion in *wspA* at ΔQ287, suggesting this second mutation is likely compensatory, allowing the Δ*lasR* RhlI R41K strain revert to the LCV state. While these exact mutations in *wspA* have not been characterized to our knowledge in SCV formation, similar compensatory mutations of *wspA* in WT strains have been associated with conversion and reversion of the SCV state [60]. Together, these data demonstrate that QS status can govern the timing and magnitude of SCV formation, and that mutations in *lasR, rhlI* and *wsp* can promote this phenotype.

To characterize the properties of the SCV subpopulation, we examined biofilm formation and antibiotic tolerance. Previous work established that SCVs derived in different genetic backgrounds and growth conditions from those under investigation here exhibited enhanced biofilm formation and antibiotic tolerance [61]. SCVs and LCVs (both Δ*lasR* RhlI R41K genotype, isolated from the same plate) were grown on Congo red agar for five days. LCVs formed biofilms with rugose centers, a pitted ring, and a smooth periphery (**Figure 1D**), while SCVs formed biofilms with hyper-rugose colonies with rugosity extending to the periphery of the colony (**Figure 1E**). Thus, we hypothesized that SCVs produce more biofilm matrix components than their LCV counterparts. Next, we performed quantitative crystal violet assays to determine if SCVs produced significantly more biofilm matrix. Indeed, SCV-forming strains in a Δ*lasR* or Δ*lasR* RhlI R41K background produced approximately 6.7- and 20.8-fold more matrix components than their isogenic LCV-forming strains (**Figure 1F**). We also tested biofilm formation of SCVs under polymyxin treatment and observed that SCV biofilms were more recalcitrant to polymyxin than biofilms formed by LCV strains (**Supplementary Figure 1C**), as others have previously reported [57, 62]. These data establish that the QS-associated SCV subpopulations identified here are phenotypically consistent with previously described *P. aeruginosa* SCV populations and genotypes [7, 8, 11].

### Transcriptomic profiling of SCVs reveals upregulation of the PhoPQ two-component system

To determine the effects of SCV formation on gene expression, we performed RNA-seq on an SCV isolate from a Δ*lasR* RhlI R41K strain and compared it to its LCV counterpart from the same plate. SCV formation in this strain typically occurs within 24 hours in rich media; thus, both LCV and SCV monocultures were grown in FDS minimal media for 24 hours prior to RNA extraction to slow differentiation and ensure stable, isogenic comparisons. Consistent with the enhanced biofilm formation described above, upregulation of *fim*, *psl*, and *pil* genes was observed (**Supplementary Table 2**). Most notable among the differentially expressed genes was the significant upregulation of the *phoPQ* operon in SCVs, with log_2_-fold-changes of 3.4 and 3.8 for *phoP* and *phoQ*, respectively (**Figure 2A**; **Supplementary Table 2**). The PhoPQ TCS has previously been linked to the formation of SCV subpopulations [61], the development of polymyxin resistance [62], and the regulation of virulence [57, 59] in several bacterial genera, consistent with the phenotypic properties of SCVs from **Figure 1**. Outstanding to us was that the QS genes *rhlI* and *rhlR* were downregulated (log_2_-fold-changes of 1.4 and 2.4, respectively) as well as other QS-regulated genes (**Supplementary Table 2**). This is consistent with the observed hyper-rugose biofilm formation observed in *rhlR* deletion strains [26]. These transcriptomic data therefore implicate the PhoPQ system as a potential regulator of both SCV formation and QS signaling in Δ*lasR* strains.

**Figure 2.**
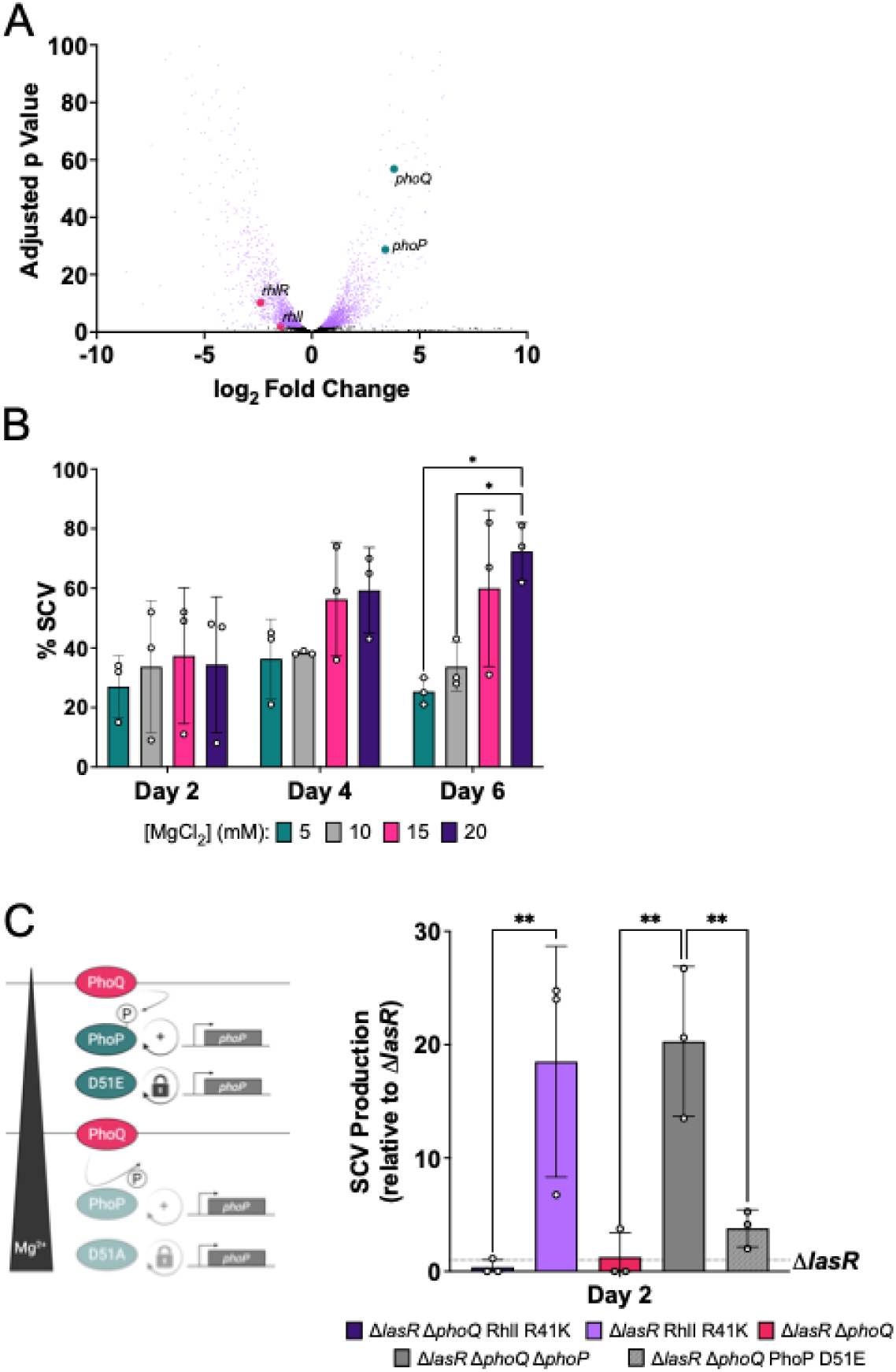
Differential regulation of the PhoPQ system in SCV populations. (**A**) Volcano plot of RNA-seq data comparing gene expression in SCV isolates relative to LCV isolates from the same plate. Selected differentially expressed loci are labeled: *phoPQ* operon (*phoP* log_2_FC = +3.4; *phoQ* log_2_FC = +3.8) and *rhlI* and *rhlR* (*rhlI* log_2_FC = −1.4; *rhlR* log_2_FC = −2.4). All differentially expressed genes in the SCV-LCV comparison can be found in **Supplementary Table 2**. (**B**) SCV production by a Δ*lasR* RhlI R41K strain passaged in FDS defined minimal media with 5 mM or 20 mM MgCl_2_ over 7 days. (**C**) (Left) Schematic of the PhoPQ response to different Mg^2+^ concentrations. At low Mg^2+^ levels, PhoP is phosphorylated and activates its own transcription in a positive-feedback loop, and at high Mg^2+^ levels PhoP is dephosphorylated, and the system is turned off. The D51E PhoP variant is a phosphor-mimic and would result in constitutive expression of *phoP* while a D51A variant is unable to be phosphorylated and would result in constitutive inactivation. (Right) SCV formation on day 2 of passaging in LB, normalized to Δ*lasR* (y = 1). Error bars represent standard deviations of biological replicates. Statistical analyses were performed using a two-way ANOVA with a Tukey’s multiple comparisons tests. P values: ns ≥0.05; * <0.05; ** <0.01; *** <0.001; **** <0.0001.

### PhoPQ contributes to SCV formation in a magnesium-dependent manner

Canonically, the PhoPQ system responds to extracellular Mg^2+^ concentrations in nearly all bacterial species for which it has been studied, including *P. aeruginosa* [57]. To determine whether PhoPQ activity contributes to SCV formation, we passaged the Δ*lasR* RhlI R41K strain in a defined minimal media supplemented with 5, 10, 15, and 20 mM of MgCl_2_ (**Figure 2B**). We note that in this minimally defined media, concentrations of MgCl_2_ below 1 mM were not tolerated for growth. Thus, we did not anticipate eliminating or maximizing SCV formation under these conditions. However, over the course of the 6-day passaging, we observed a stratification of SCV formation as a function of Mg^2+^ concentration; specifically, higher concentrations of MgCl_2_ resulted in higher proportions of SCVs by day 6 (**Figure 2B**). To test if lower levels of Mg^2+^ could further suppress SCV formation, we used standard LB rich media. LB media is undefined in its Mg^2+^ concentration, but direct measurements of its chemical composition reported an approximate final concentration of 100 µM [63]. We next added an additional 25 µM or 5 mM of MgCl_2_ to LB media and assayed for SCV formation in either a Δ*lasR* RhlI R41K strain or a Δ*lasR*Δ*phoQ* RhlI R41K strain. We expected that the strain lacking *phoQ* would be unresponsive to fluctuations in Mg^2+^ and, indeed, that was the case (**Supplementary Figure 2A**). Overall, 5 mM MgCl_2_ maintained moderate levels of SCV formation (∼20%) relative to 25 µM MgCl_2_ in both passaging experiments performed in LB and defined minimal media. Growth of different strains were comparable in each media type (**Supplementary Figure 2B and 2C**).

To further define the genetic components underpinning SCV formation, we performed a 6-day passaging experiment in LB media without exogenous Mg^2+^ supplementation. We chose this condition and timing component because this is when we observed enhanced SCV formation in a QS-dependent manner (**Figure 1C**). The data in Figure 2C are normalized to the percentage of SCV formed on day 2 by a Δ*lasR* strain (average = 2.6%, N=12). In a Δ*lasR* background with WT RhlI or RhlI R41K, we deleted *phoQ* and assessed SCV formation (**Figure 2C**). The deletion of *phoQ* in both the Δ*lasR* and Δ*lasR* RhlI R41K strains resulted in a nearly complete loss of SCV formation on day 2 (**Figure 2C**), indicating that the unphosphorylated state of PhoP is repressive of SCV formation. This trend continued into day 4 for the Δ*phoQ* Δ*lasR* RhlI R41K strain (**Supplementary Figure 2D**). Conversely, when *phoP* was deleted in a strain lacking *lasR* and *phoQ*, SCV formation was restored (**Figure 2C**), indicating that the observed repression phenotype was specifically through PhoP function. We note that we were unable to make single clean deletions of *phoP* alone, limiting our genetic analyses to the strains shown in **Figure 2C**.

We hypothesized that the strain expressing a PhoP D51E phospho-mimic variant would have a phenotype similar to the deletion of *phoP* in a Δ*lasR*Δ*phoQ* background. Interestingly, introducing a PhoP D51E variant on the chromosome in a Δ*lasR*Δ*phoQ* strain resulted in only a slight increase in SCV formation (**Figure 2C**). However, PhoP is known for its auto-regulatory positive feedback loop, and we suspect that *phoP* expression levels play an important role in regulating SCV formation. Altogether, these data suggest that the PhoPQ system co-regulates SCV formation with QS state, particularly the Rhl system.

### PhoPQ co-regulates RhlR-dependent virulence factor production in the absence of LasR

Given that *rhlIR* and *phoPQ* were differentially expressed in opposite directions in an SCV dependent manner, and the role of the PhoPQ system in regulating SCV production (**Figure 2**), we next examined the role of the PhoPQ system on RhlR-dependent traits. We used pyocyanin production as a readout because phenazine gene expression is regulated by RhlR-dependent signaling. We deleted *phoQ* from WT and SCV-forming strains and grew them in FDS minimal media with time points collected at 5 and 18 hours. The Δ*phoQ* single deletion strain was moderately delayed (*p=*0.1338) in pyocyanin production at 5 hours compared to WT; this deficiency was corrected by the 18-hour timepoint and WT and Δ*phoQ* strains produced nearly identical amounts of pyocyanin (**Figure 3A**). Meanwhile, the Δ*lasR*Δ*phoQ* strain produced pyocyanin at levels comparable to the Δ*lasR* parent strain at both timepoints (**Figure 3A**). In contrast, deletion of *phoQ* in the Δ*lasR* RhlI R41K background significantly reduced the elevated pyocyanin levels characteristic of that strain at both the 5-hour and 18-hour time points (**Figure 3A**), indicating that PhoQ-dependent phosphorylation of PhoP is required for full pyocyanin production in this background and under these specific growth conditions.

**Figure 3.**
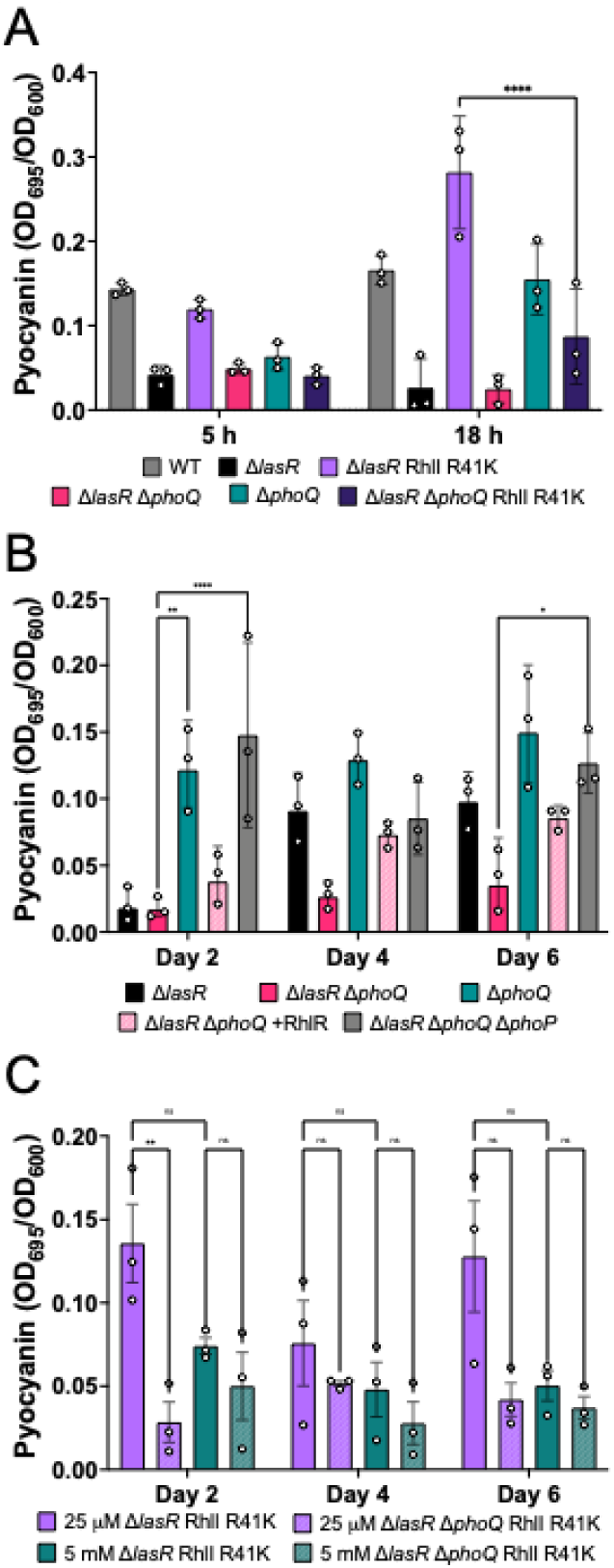
The role of the PhoPQ system in RhlR-dependent pyocyanin production. (**A**) Pyocyanin production of Δ*phoQ*, Δ*lasR*Δ*phoQ*, and Δ*lasR*Δ*phoQ* RhlI R41K strains in FDS minimal media over 18 hours. Values for WT, Δ*lasR*, and Δ*lasR* RhlI R41K are from Figure 1A and are provided to act as a direct comparison to their otherwise isogenic *phoQ* deletion counterparts. All experiments were performed at the same with appropriate controls. (**B**) Longitudinal pyocyanin production of the same set of strains from Figure 2C in LB over 6 days. Additionally, *rhlR* overexpression in Δ*lasR*Δ*phoQ* and Δ*lasR*Δ*phoQ* RhlI R41K strains are shown. (**C**) Longitudinal pyocyanin production for Δ*lasR* RhlI R41K (solid bars) or Δ*lasR*Δ*phoQ* RhlI R41K (dashed bars) in LB rich media with 25 µM (purple) or 5 mM (teal) MgCl_2_. Error bars represent standard deviations of biological replicates. Statistical analyses were performed using a two-way ANOVA with a Tukey’s multiple comparisons tests. P values: ns ≥0.05; * <0.05; ** <0.01; *** <0.001; **** <0.0001.

To assess the effects of *phoQ* deletion on QS-dependent phenotypes on a time scale like that of SCV formation, we monitored pyocyanin production for six days of serial passaging in LB. To correlate population-wide behaviors such as SCV formation and pyocyanin production, we used the same strains as shown in **Figure 2C**. The Δ*phoQ* single deletion strain maintained relatively high levels of pyocyanin throughout time course (**Figure 3B**), consistent with the observed pyocyanin production on a shorter time scale (**Figure 3A**). Meanwhile, the Δ*lasR* strain recovered pyocyanin production after day four (**Figure 3B**), at the onset of when the isolate initiates SCV formation (**Figure 2C**). Interestingly, a Δ*lasR*Δ*phoQ* strain exhibited minimal pyocyanin production across all days compared to the Δ*lasR* strain, indicating that, over time,

PhoQ plays a role in regulating RhlR-dependent traits when strains exhibit a loss of LasR signaling (**Figure 3B**). A similar trend was observed for the comparison between Δ*lasR*Δ*phoQ* RhlI R41K and Δ*lasR* RhlI R41K (**Figure 3A**), further supporting this notion, although the differences in pyocyanin production were not statistically significant over the time-course in LB (**Supplementary Figure 2E**). We hypothesize that this is because the RhlI R41K variant provides a buffering effect for the loss of *phoQ* in the Δ*lasR* background.

The loss of pyocyanin production in a Δ*lasR*Δ*phoQ* strain compared to a Δ*lasR* strain supported a potential mechanism where unphosphorylated PhoP represses pyocyanin production. To test whether PhoP was specifically required for this repression, we deleted *phoP* in the Δ*lasR*Δ*phoQ* strain and measured pyocyanin production. Indeed, repression of pyocyanin was relieved in the triple mutant during the earliest timepoint (**Figure 3B**), indicating that PhoP likely represses at least one component of Rhl signaling cascade or its downstream regulation of phenazine gene expression (**Supplementary Figure 3A**). To confirm that the pyocyanin deficiency in a Δ*lasR*Δ*phoQ* strain was attributable to reduced RhlR activity, we introduced a second copy of *rhlR* on an IPTG-inducible plasmid. Overexpression of *rhlR* partially complemented pyocyanin production (**Figure 3B**). Overall, the lack of pyocyanin production (**Figure 3B**) correlated with the decline of SCV formation (**Figure 2C**) in LB on the time scale tested.

To further define the role of PhoPQ signaling in regulating QS behaviors, we performed a pyocyanin production time course assay for six days in a Δ*lasR* R41K background with or without *phoQ* in the presence of either 25 µM or 5 mM MgCl_2_ in LB, similar to the experiments performed in **Supplemental Figure 2** for SCV formation. Low levels of magnesium drove pyocyanin production in a PhoQ-dependent manner on day two (**Figure 3C**); specifically, the Δ*lasR* RhlI R41K produced high levels of pyocyanin relative to the Δ*lasR*Δ*phoQ* RhlI R41K at the 25 µM condition, indicating that the flow of phosphoryl groups through the PhoPQ system is important for expressing the phenazines via RhlR-dependent signaling.

### Unphosphorylated PhoP directly represses the *rhlR* promoter

Based on the loss of pyocyanin production in a Δ*lasR* Δ*phoQ* RhlI R41K strain relative to a Δ*lasR* RhlI R41K strain (**Figure 3A**), we hypothesized that the activity of PhoQ on PhoP results in the repression of *rhlR*. Given that PhoQ is not present to phosphorylate PhoP, we hypothesized that the unphosphorylated state of PhoP represents the repressive form of PhoP for the *rhlR* promoter. Further supporting this hypothesis, the RNA-seq experiment (**Figure 2A**) that revealed increased *phoPQ* expression with a concomitant decrease in *rhlIR* expression was performed in a media containing high levels of Mg^2+^, which would drive the PhoP population into the unphosphorylated state. Additionally, a previous ChIP-seq study demonstrated that PhoP bound to the *rhlR* promoter [57]. Given our inability to delete *phoP* in all the relevant genetic backgrounds and to explicitly test this in isolation, we constructed a three-plasmid *Escherichia coli* reporter system. We co-expressed *lasR* from an arabinose-inducible promoter (pBAD) to serve as a positive activator of p*rhlR*-*luxCDABE* alongside *phoP* expressed from an IPTG-inducible promoter (pACYC). As expected, expression of LasR alongside an empty pACYC plasmid in the presence of the p*rhlR* reporter resulted in high levels of light production (**Figure 4A**). Additionally, when the converse experiment was performed with only *phoP* expressed from pACYC alongside an empty pBAD plasmid, no light was produced from the p*rhlR* reporter. Co-expression of LasR and PhoP resulted in a significant reduction in light production from the pBAD-*lasR* only control, indicating that PhoP likely competes with LasR at the *rhlR* promoter to result in inactivation of the reporter (**Figure 4A**). We were unable to construct a strain that expressed the *phoQ* gene from *P. aeruginosa,* nor could we perform our light assay in minimal media with the low levels of Mg^2+^ required to drive PhoQ phosphorylation of PhoP. Thus, to probe the importance of PhoP phosphorylation state, we introduced two variants on the pACYC-*phoP* plasmid that resulted in a locked state: PhoP D51A (unable to be phosphorylated) and PhoP D51E (phosphomimic). Consistent with the loss of *phoQ* in *P.* aeruginosa, which led to unphosphorylated PhoP and, thus, the repression of *rhlR* expression, the PhoP D51A variant repressed p*rhlR*-driven luminescence to a level comparable to WT PhoP (**Figure 4A**). Conversely, the PhoP D51E variant produced luminescence indistinguishable from the empty vector control, failing to repress the promoter (**Figure 4A**). Thus, the *E. coli* reporter system is consistent with our *in vivo* interpretation of PhoQ-dependent regulation of pyocyanin production via diminished *rhlR* expression.

**Figure 4.**
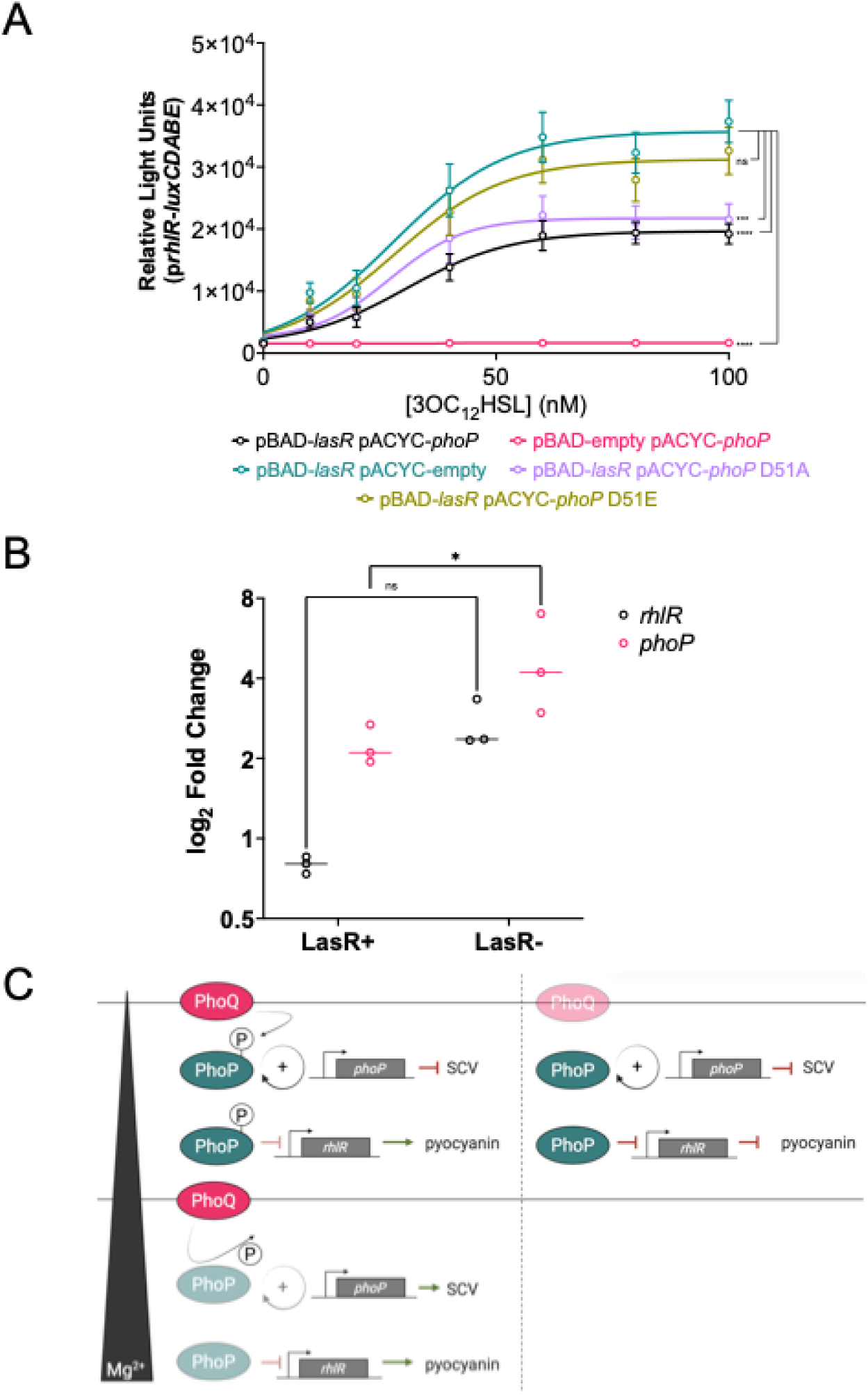
PhoP is a repressor of *rhlR.* (**A)** Luciferase assay performed in *E. coli* showing relative light units produced from the *rhlR* promoter (pCS26) under regulation of LasR (pBAD) and PhoP (pACYC). (**B**) Transcript levels of *rhlR* and *phoP* in phosphorylation-locked variant PhoP D51E relative to PhoP D51A in wild-type (LasR+) and Δ*lasR* (LasR-) backgrounds. Statistical analyses were performed using a two-way ANOVA. *P* values: ns ≥ 0.05; * <0.05; ** <0.01; *** <0.001; **** <0.0001. (**C**) Schematic of PhoPQ regulation of *rhlR* and downstream effects on pyocyanin and SCV production.

To further validate the *E. coli* reporter system *in vivo*, we performed qRT-PCR on *rhlR* in *P. aeruginosa* strains expressing *phoP* D51E relative to strains expressing *phoP* D51A in WT or Δ*lasR* backgrounds. Consistent with the autoregulatory feedback of phosphorylated PhoP, *phoP* levels were elevated in the strain expressing *phoP* D51E relative to *phoP* D51A, regardless of strain background. Conversely, *rhlR* transcription increased in the strain with constitutively active PhoP lacking *lasR* (**Figure 4B**). Interestingly, *phoP* levels were also significantly increased in this same background, indicating that the initial loss of LasR function is key to driving this shift in transcriptional reprogramming that alters QS and, perhaps, by proxy SCV formation. This trend was consistent across genotypes (**Supplementary Figure 3B**) lacking *lasR*. These data are consistent with previous findings that showed that disrupting QS, specifically RhlR-dependent signaling, can result in elevated expression of *phoPQ* via an unknown mechanism [64, 65]. Collectively, these data support a model in which unphosphorylated PhoP binds the *rhlR* promoter to repress QS signaling, and PhoQ-dependent phosphorylation of PhoP relieves this repression to de-repress *rhlR* and activate downstream virulence factor production (**Figure 4C**).

## DISCUSSION

SCV formation is an adaptive stress response conserved in several bacterial genera. Other groups have demonstrated that *P. aeruginosa* SCVs are induced by the accumulation of stable mutations [7, 9, 11]. Here, we show that SCV formation is promoted by *lasR* and *rhlI* mutations, and that the PhoPQ system is an upstream regulator of SCV formation and pyocyanin production in a Δ*lasR* background. The regulation of SCVs is likely indirect through unknown mechanisms whereas pyocyanin expression is through direct regulation of RhlR in the absence of *lasR.* We showed previously that *rhlI* mutations can enhance virulence phenotype expression in a Δ*lasR* background, and this was also true for the RhlI R41K mutation we identified in a *P. aeruginosa* isolate obtained from a patient with cystic fibrosis. The Δ*lasR* RhlI R41K strain is a hyper-producer of both pyocyanin and SCVs, demonstrating how naturally evolved mutations in QS pathways can enhance virulence of clinical strains. Moreover, the observation that production of SCVs is enhanced in Δ*lasR* strains could partially explain the high frequency of *lasR* mutations reported in clinical strains, especially if SCVs are better adapted to the infection niche.

Other groups have shown that the transcription factor PhoB of the PhoBR TCS regulates *rhlR* expression under phosphate limiting conditions [48, 49, 66]. We identified *phoPQ,* another TCS, in our SCV RNA-seq dataset (**Figure 2A**). Deletion of *phoQ* delayed, but did not abrogate, virulence phenotype expression in Δ*lasR* strains (**Figure 3**) and we observe over-production of SCVs (**Figure 2C**) and pyocyanin (**Figure 3B**) in a Δ*lasR*Δ*phoQ*Δ*phoP* strain, suggesting a repressive role of PhoP. We further show that unphosphorylated PhoP represses the *rhlR* promoter using an *in vitro* luciferase reporter assay and qRT-PCR (**Figure 4A** and **4B**), which corroborates a previous ChIP-seq report showing that PhoP has a binding site in the *rhlR* promoter [61]. We note that we were unable to make a *phoP* single deletion in a Δ*lasR* background, which could be an indication of its essentiality in this context. This is the first report, to our knowledge, of direct QS regulation by the PhoPQ system. This regulation is likely part of PhoPQ nutrient response signaling, as different concentrations of magnesium affected the proportions of SCV formation and pyocyanin production in a Δ*lasR* RhlI R41K strain (**Figures 2B and 3C**). The finding that magnesium sensing alters QS states is biologically relevant to the infection niche in the human lung, where magnesium modulates mucus clearance and inflammation [67, 68]. In the case of cystic fibrosis patients, CFTR mutations result in magnesium deficiency and are often treated with oral magnesium supplementation [69]. Based on the data presented here, variations in magnesium levels in the cystic fibrosis airway might influence SCV production of *P. aeruginosa* in the lung niche.

Our data support a model where, under high magnesium concentrations, PhoQ dephosphorylates PhoP and causes repression of QS, which results in the loss of SCVs and pyocyanin production. In *Salmonella typhimurium*, which has a homologous PhoP protein to that in *P. aeruginosa,* Mg^2+^ levels in the low millimolar range resulted in unphosphorylated PhoP and repression of PhoP-regulated genes, whereas concentrations of Mg^2+^ in the micromolar range increased PhoP phosphorylation and target gene expression [70, 71]. However, evidence of PhoQ acting as both a kinase under low magnesium conditions and a phosphatase under high magnesium conditions in *P. aeruginosa* has been previously reported [52] and, based on the data we present here, it is the kinase activity of PhoQ and de-repression of QS in a Δ*lasR* strain that is important for the SCV phenotype. This suggests that phosphorylation of PhoP, rather than its dephosphorylation, is the mechanism by which PhoP target genes are de-repressed in *P. aeruginosa*.

In *S. typhimurium*, PhoP positively activates its own transcription when phosphorylated under low magnesium concentration [72], but our data in *P. aeruginosa* shows that *phoP* is significantly upregulated in Δ*phoQ* strains (**Supplementary Figure 3**). We also show that increasing concentrations of Mg^2+^ in the millimolar range caused an increase in SCV production (**Figure 2B**) which is seemingly counter-intuitive to our model. However, the expression of *phoP* itself would be repressed under high magnesium concentrations; therefore, we interpret the increasing production of SCVs under high Mg^2+^ to be a result of low PhoP levels in the cell due to the autoregulatory mechanism of the PhoPQ system and the subsequent lack of repression on *rhlR.* Indeed, the significant increase in *phoP* expression in Δ*phoQ* deletion strains (**Supplementary Figure 3**) correlate with the loss of virulence phenotypes in Δ*lasR* backgrounds while further deletion of *phoP* causes over-expression of these phenotypes (**Figures 2C and 3B**). Furthermore, the inability of PhoP D51E to completely de-repress SCV production in a Δ*phoQ* background also suggests that the levels of PhoP, rather than its phosphorylation state, dictate the level of regulation at the *rhlR* promoter (**Figures 2C and 4C**) because constitutive phosphorylation would increase the levels of PhoP itself. Moreover, our data provide further evidence that virulence factor expression converges on RhlR regulation and activation in *P. aeruginosa*. Importantly, none of the genetic manipulations we made in this study completely abolished SCV production, suggesting that there are other regulators that help coordinate SCV differentiation, potentially systems that are generally repressed by LasR.

## METHODS

### Strain and plasmid construction

Gene mutation and deletion constructs were cloned into a pEXG2 suicide vector using NEB HiFi assembly. After confirming sequences, plasmids were transformed into *E. coli* SM10 conjugative competent cells by heat-shock. Matings were performed by co-streaking *P. aeruginosa* and *E. coli* on LB plates and incubating statically at 37 °C overnight. Mixed culture patches were scraped from plates, washed in 1 mL LB, pelleted, resuspended in 100 μL LB, plated on 30 μg/mL gentamicin plates, and incubated at 37 °C overnight. Individual colonies were recovered in 1 mL LB at 37 °C for 2 h. Cultures were then pelleted, resuspended in 100 μL LB, plated on 5% sucrose plates, and incubated at 30 °C overnight. Individual colonies were dual selected on 30 μg/mL gentamicin and LB plates. Colonies that grew on LB but not on gentamicin were submitted for sequencing to confirm chromosomal recombination. The pUCP18-p*lac*-*rhlR* plasmid was miniprepped from an *E. coli* glycerol stock and electroporated into glycerol-competent *P. aeruginosa* strains. Transformants were selected on carbenicillin (400 μg/mL) plates. A complete list of primers, plasmids, and strains is provided in **Supplementary Table 3**.

### Whole genome sequencing

Genomic DNA was extracted using the PacBio Nanobind CBB kit following the “extracting HMW DNA from Gram-negative bacteria” protocol. Briefly, single colonies were grown overnight in 3 mL LB at 37°C with shaking; 400 μL of culture was used to achieve a cell count within the target range of 5 x 10^8^ to 5 x 10^9^ cells. Extracted DNA was quantified by NanoDrop and sequenced on an Oxford Nanopore platform. Reads were trimmed using PoreChop v0.2.4. *De novo* assemblies were generated using Flye v2.9.4 [73], and unique SNPs were identified using Snippy 4.6.0 [74].

### Pyocyanin and SCV assays in minimal media

Strains were cultured from freezer stocks in 3 mL LB overnight at 37°C with shaking. High-cell density cultures (OD_600 nm_ ≈ 2.0) were back-diluted 1:10 into 5 mL FDS media (2% glycerol, 10 g/L DL-alanine, 50 μM iron (III) citrate, 0.1 M Na_2_SO_4_, 20 mM MgCl_2_, 500 μM K_2_HPO_4_) and incubated for 18 hours. Time points were collected at 5 and 18 hours: 1 mL of culture was removed, and growth was measured at OD_600 nm_ following a 1:10 dilution in FDS media. The remainder was pelleted and the absorbance of cell-free supernatant was measured at OD_695_ to quantify pyocyanin. For the SCV magnesium dependence assay, the Δ*lasR* RhlI R41K strain was passaged in FDS media with either 5 mM or 20 mM MgCl_2_ for one week; cultures were serially diluted 1:10 in LB and the 10^−7^ dilution was plated on LB agar to enumerate colonies.

### Longitudinal pyocyanin and SCV assays

Strains were cultured from freezer stocks in 3 mL LB overnight at 37°C with shaking. At 24 hours, 1 mL of culture was collected; growth was measured at OD_600_ nm following a 1:10 dilution in LB, and the remainder was pelleted. The absorbance of cell-free supernatant was measured at OD_695 nm_ to quantify pyocyanin, and cell pellets were reserved for RNA extraction. Strains were then diluted 1:100 into 3 mL fresh LB and grown for 24 h. This process was repeated every 24 h over the course of one week.

### Microscopy

Plates were imaged using an Olympus SZX12 Stereo Fluorescence Microscope equipped with a Nikon DS-Fi3 camera and NIS Elements software. Biofilm plates were imaged at 7x magnification and colony images were captured at 12.5x magnification.

### Crystal violet assays

Overnight cultures were diluted 1:100 into different concentrations of polymyxin B, and 200 μL of each strain was dispensed in triplicate into a 96-well plate. Plates were incubated statically at 37 °C overnight. After 24 hours, planktonic bacteria were transferred to a separate 96-well plate, and OD_600 nm_ was measured. Remaining biofilms were stained with 1% crystal violet and solubilized with acetic acid; crystal violet staining was quantified at OD_560_ using a GloMax plate reader (Promega).

### Minimum inhibitory concentration determination

Overnight cultures were grown at 37 °C with shaking at 200 rpm. The following day, cultures were diluted 1:20 (100 μL into 2 mL LB) and incubated at 37 °C for 2 to 2.5 h until OD_600 nm_ was between 0.4 and 0.6. Each well of a 96-well plate received 90 μL LB followed by 10 μL of back-diluted culture. A 30 mg/mL polymyxin B stock was serially diluted 1:100,000 in sterile water and added to wells to achieve final concentrations ranging from 0 to 3.25 mg/mL. Plates were incubated overnight at 37 °C with shaking at 350 rpm, and OD_600 nm_ was measured the following day. Experiments were performed in triplicate, and data were analyzed using GraphPad Prism. The sub-MIC concentration for polymyxin B was determined to be 0.001625 mg/mL.

### Sub-inhibitory antibiotic induction of SCV formation

Overnight cultures were grown at 37 °C with shaking at 200 rpm and diluted 1:10 the following day (300 μL into 3 mL LB). One untreated and three treated samples were prepared per strain. Based on the sub-MIC concentration of polymyxin B (0.001625 mg/mL), treated samples received 16 μL of a 1:100 dilution of a 30 mg/mL polymyxin B stock. Cultures were incubated statically at 37 °C for three days, after which 100 μL of each culture was serially diluted to 10^−7^. The 10^−4^ through 10^−7^ dilutions were plated on LB agar and spread with beads. Plates were incubated at 37 °C overnight and SCV counts were determined the following day. Experiments were performed in triplicate, and data were analyzed using GraphPad Prism.

### RNA-seq and data analysis

The Δ*lasR* RhlI R41K strain was passaged in LB for one week and individual colonies were picked from the same serial dilution plate. SCV and LCV isolates were grown in separate 3 mL monocultures in LB overnight at 37 °C with shaking. RNA was extracted using a phenol-chloroform method: pelleted cultures were resuspended in 500 μL TRIzol reagent and transferred to screw-cap tubes containing 100 μL zirconia beads. Samples were homogenized using a bead-beater at 4 °C for 2 min. Chloroform (100 μL) was added, samples were briefly shaken and incubated at room temperature for 2 minutes, then centrifuged at maximum speed on a chilled centrifuge for 5 minutes. The aqueous layer was transferred to a new tube, 1 mL isopropanol was added, and samples were inverted to precipitate RNA and centrifuged for 1 minute at maximum speed. The isopropanol was aspirated; samples were washed with 1 mL of 80% ethanol, centrifuged, and the ethanol was allowed to evaporate before RNA was resuspended in nuclease-free water. RNA was quantified by NanoDrop and treated with the TURBO DNase kit per manufacturer instructions (Thermo Fisher Scientific). cDNA libraries were constructed using the NEB Next Ultra II RNA Library Preparation Kit for Illumina per manufacturer instructions. Libraries were resuspended in 20 μL nuclease-free water and analyzed by TapeStation. Sequencing was performed on an Illumina MiSeq platform. Reads were mapped to the *P. aeruginosa* UCBPP-PA14 genome (NCBI accession number NC_008463.1) using the BWA-MEM algorithm within SAMtools v1.1.0. Differential expression analysis was performed using DESeq2 (v1.52.0) [75]. RNA-Seq data is accessible at the NCBI SRA BioProject PRJNA1465858.

### qRT-PCR

Strains were cultured from freezer stocks in 3 mL LB overnight at 37 °C with shaking. High-cell density cultures (OD_600 nm_ ≈ 2.0) were back-diluted 1:100 into 3 mL FDS media and incubated for 18 hours. Strains were then passaged twice more (1:100 back-dilutions in fresh FDS media) for a total of three days. Total cellular RNA from 3 mL passage-2 cultures was isolated using the PureLink RNA Mini Kit (Invitrogen). RNA was quantified by NanoDrop and treated with the TURBO DNase kit per manufacturer instructions (Thermo Fisher Scientific). cDNA was synthesized using the SuperScript III Reverse Transcriptase kit (Invitrogen) in 20 μL total reaction volumes. Each qPCR reaction contained 2× SYBR Select Master Mix (Applied Biosystems), 200 nM primers, and 2 μL of 1:5-diluted cDNA in a total volume of 20 μL. Amplification and cycle threshold quantification were performed on a 7500 Fast Real-Time PCR system (Applied Biosystems) using software v2.3. The 16S rRNA gene was used as a housekeeping reference.

### *E. coli* light assay

The three-plasmid system used for luciferase reporter experiments comprises pCS26-p*rhlR*-*luxCDABE*, pBAD-*lasR*, and pACYC-*phoP*, along with their respective empty vector controls. Plasmids were co-transformed into BL21 calcium-competent *E. coli* cells. Strains were grown overnight in LB with appropriate antibiotics at 37 °C with shaking, then back-diluted 1:1000 in antibiotic-containing media until OD_600_ reached 0.4 to 0.6. Cultures were dispensed into a 96-well plate and supplemented with 0.1% arabinose, 0.01 mM IPTG, and 3OC_12_HSL to induce expression from the pBAD and pACYC plasmids, respectively. Plates were incubated for 4 h at 37 °C with shaking, and luminescence was measured on a GloMax plate reader (Promega).

## ACKNOWLEDGMENTS

The authors thank all members of the Paczkowski laboratory and the research groups in the Division of Genetics at the Wadsworth Center for helpful discussions. This work was made possible by the dedicated staff scientists at the Advanced Genomics Technologies Cluster and the Media & Tissue Core facilities at the Wadsworth Center.

