## Supplemental Figures for "PhoPQ regulates quorum sensing in small colony variant subpopulations of *Pseudomonas aeruginosa*"

**SUPPLEMENTARY MATERIALS**

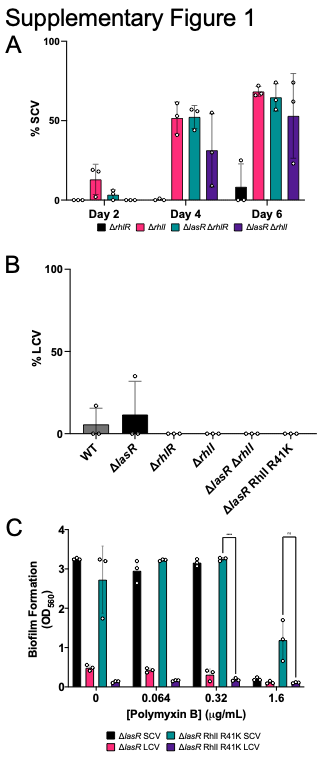

**Supplementary Figure 1.** QS mutants produce a stable SCV morphology over time. (**A**) Small colony percentages after passaging strains in LB for 6 days. (**B**) Percentage of revertant colonies after passaging SCV-differentiated strains for one week in LB. (**C**) Crystal violet assays were performed to quantify the biofilm formation of SCVs and LCVs in the presence of polymyxin B (μg/ML). Statistical analyses were performed using a two-way ANOVA with a Tukey’s multiple comparisons tests. P values: ns ≥0.05; * <0.05; ** <0.01; *** <0.001; **** <0.0001.

Error bars represent standard deviations of three biological replicates.

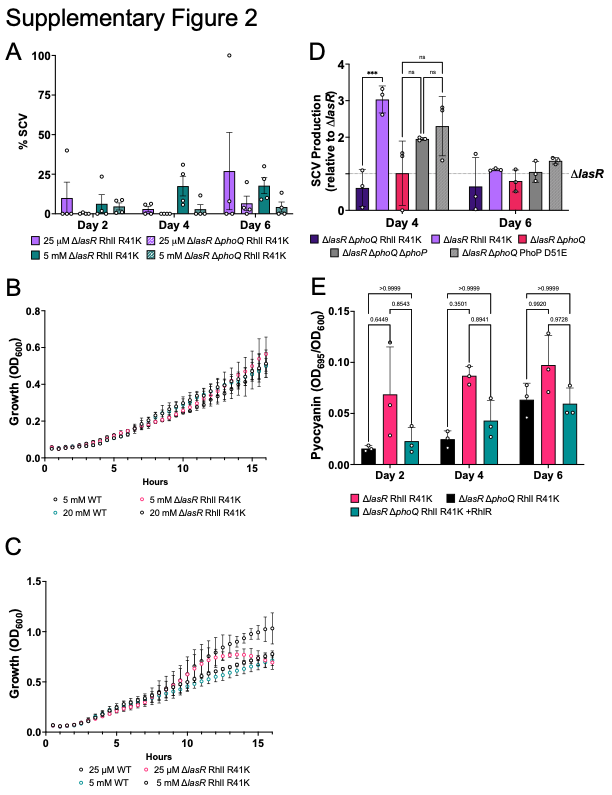

**Supplementary Figure 2.** Magnesium supports SCV formation. (**A**) Percentage of SCVs for Δ*lasR* RhlI R41K (solid) or Δ*lasR*Δ*phoQ* RhlI R41K (hash marks) grown in LB supplemented with 25 µM (purple) or 5 mM (blue) MgCl_2_. (**B**) Growth curves of WT and Δ*lasR* RhlI R41K in FDS supplemented with 5 or 20 mM MgCl_2_ and (**C**) in LB supplemented with 25$\mu$M or 5mM MgCl_2_. (**D**) SCV formation on days 4 and 6 of passaging in LB, normalized to ∆*lasR* (y = 1). (**E**) Pyocyanin production of Δ*lasR* RhlI R41K, Δ*lasR* Δ*phoQ* RhlI R41K and Δ*lasR* Δ*phoQ* RhlI R41K + pJN105-*rhlR* on days 2, 4 and 6 of passaging in LB. Error bars represent standard deviations of three biological replicates. Statistical analyses were performed using a two-way ANOVA with a Tukey’s multiple comparisons tests. P values: ns ≥0.05; * <0.05; ** <0.01; *** <0.001; **** <0.0001.

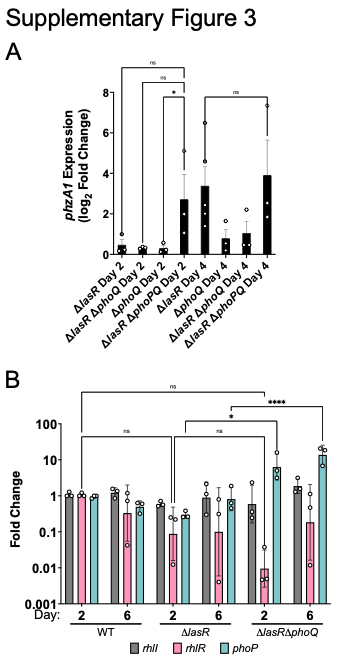

**Supplementary Figure 3.** Temporal transcription of quorum sensing genes in SCV producing strains. (**A**) qRT-PCR of *phzA1* in different strain backgrounds on days 2 and 4 of passaging in LB media. (**B**) qRT-PCR data for *rhlI*, *rhlR*, and *phoP* in mixed-phenotype strains on days 2 and 6 of passaging, normalized to WT day 2 cultures. Error bars represent standard deviations of three biological replicates. Statistical analyses were performed using a two-way ANOVA. *P* values: ns $\geq$ 0.05; * $<$0.05; ** $<$0.01; *** $<$0.001; **** $<$0.0001.

**Supplementary Table 1.** Mutated loci observed in whole genome sequencing of SCV and SCV revertant isolates.

| Strains with variant  (SC=small colony) | Type | Ref  Seq | Alt Seq | Genome Annotation | Loci | Amino Acid Position | Effect |
| --- | --- | --- | --- | --- | --- | --- | --- |
| Δ*lasR* RhlI (R41K) Revert 4 | snp | A | G | RlpA family lipoprotein | PA14_12090 | 255/341 | Synonymous variant |
| Δ*lasR* SC 1  Δ*lasR* SC 2 | snp | G | A | WspA chemotaxis transducer | PA14_16430 | 344/542 | Missense variant |
| Δ*lasR* RhlI (R41K) SC 1  Δ*lasR* RhlI (R41K) SC 2  Δ*lasR* RhlI (R41K) Revert 1 Δ*lasR* RhlI (R41K) Revert 2 Δ*lasR* RhlI (R41K) Revert 3 Δ*lasR* RhlI (R41K) Revert 4 | snp | G | A | WspA chemotaxis transducer | PA14_16430 | 464/542 | Missense variant |
| Δ*lasR* RhlI (R41K) Revert 1 Δ*lasR* RhlI (R41K) Revert 2 Δ*lasR* RhlI (R41K) Revert 3 Δ*lasR* RhlI (R41K) Revert 4 | del | AAGC | A | WspA chemotaxis transductor | PA14_16430 | 286/542 | Conservative in frame del |
| Δ*lasR* RhlI (R41K) SC 2  Δ*lasR* RhlI (R41K) Revert 1 | ins | G | GT | ZbdP | PA14_23420 | 291/722 | Frameshift variant |
| Δ*lasR* RhlI (R41K) Revert 1 Δ*lasR* RhlI (R41K) Revert 2 Δ*lasR* RhlI (R41K) Revert 3 Δ*lasR* RhlI (R41K) Revert 4 | snp | G | T | ZbdP | PA14_23420 | 570/722 | Missense variant |
| Δ*lasR* SC 2 | del | TGA | T | malonyl-CoA-ACP transacylase | PA14_25650 | 309/312 | Frameshift variant |
| Δ*lasR* RhlI (R41K) SC 2 | snp | C | A | hypothetical protein | PA14_33600 | 359/403 | Missense variant |
| Δ*lasR* SC 1  Δ*lasR* RhlI (R41K) SC 1  Δ*lasR* RhlI (R41K) SC 2  Δ*lasR* RhlI (R41K) Revert 2 | del | TCC | T | RNA polymerase ECF-subfamily sigma-70 factor | PA14_46660 | 9/246 | Frameshift variant |
| Δ*lasR* SC 2 | snp | A | C | hypothetical protein | PA14_48890 | 297/432 | Missense variant |
| Δ*lasR* RhlI (R41K) Revert 4 | del | GC | G | response regulator PirR | PA14_52250 | 132/239 | Frameshift variant |
| Δ*lasR* RhlI (R41K) Revert 2 | snp | T | G | S-adenosyl-methyltransferase MraW | PA14_57450 | 38/313 | Missense variant |

**Supplementary Table 2.** Differentially expressed genes between the SCV-LCV populations.

**Supplementary Table 3.** Strains, plasmids, and oligonucleotides used in this study.

| **Strains & Plasmids** | | | |
| --- | --- | --- | --- |
| **Strain #** | **Strain** | **Plasmid** | **Source** |
| JPS111 | *E. coli* DH5α | pBAD-*lasR* | Paczkowski et al. 2017 |
| JPS151 | PA14 Δ*rhlR* |  | Mukherjee et al. 2017 |
| JPS154 | PA14 Δ*rhlI* |  | Mukherjee et al. 2017 |
| JPS156 | PA14 Δ*lasR* |  | Mukherjee et al. 2017 |
| JPS157 | PA14 Δ*lasR* Δ*rhlR* |  | Mukherjee et al. 2017 |
| JPS222 | UCBPP-PA14 WT |  | Gift from Dr.George O’Toole |
| JPS226 | *E. coli* DH5α | pBAD |  |
| JPS227 | *E. coli* DH5α | pEXG2 |  |
| JPS247 | *E. coli* Top 10 | pUCP18 |  |
| JPS473 | *E. coli* DH5α | pUCP18-p*lac*-*rhlR* | Mukherjee et al. 2017 |
| JPS1013 | PA14 Δ*lasR* Δ*rhlI* |  |  |
| JPS1080 | *E. coli* SM10-λpiR | pEXG2-*rhlI* R41K | This study. |
| JPS1141 | PA14 Δ*lasR rhlI* R41K |  | This study. |
| JPS1142 | PA14 *rhlI* R41K |  | This study. |
| JPS 1689 | *E. coli* SM10-λpiR | pEXG2-Δ*phoQ* | This study. |
| JPS1744 | PA14 Δ*phoQ* |  | This study. |
| JPS1745 | PA14 Δ*lasR* Δ*phoQ* |  | This study. |
| JPS1773 | PA14 Δ*lasR* Δ*phoQ rhlI* R41K |  | This study. |
| JPS1810 | *E. coli* SM10-λpiR | pEXG2-*phoP* D51E | This study. |
| JPS1840 | PA14 Δ*lasR phoP* D51E |  | This study. |
| JPS1845 | *E. coli* DH5α | pCS26-p*rhlR* | This study. |
| JPS1960 | *E. coli* SM10-λpiR | pEXG2-Δ*phoQ* Δ*phoP* | This study. |
| JPS1991 | PA14 Δ*phoQ* Δ*phoP* |  | This study. |
| JPS1994 | PA14 Δ*lasR* Δ*phoQ* Δ*phoP* |  | This study. |
| JPS2411 | *E. coli* DH5α | pACYC-*phoP* | This study. |
| JPS2581 | *E. coli* SM10-λpiR | pEXG2-*phoP* D51A | This study. |
| JPS2629 | PA14 *phoP* D51A |  | This study. |
| JPS2682 | PA14 Δ*lasR phoP* D51A |  | This study. |
| JPS2684 | PA14 *phoP* D51E |  | This study. |
| JPS2502 | *E. coli* DH5α | pACYC-Duet |  |
| JPS2770 | *E. coli* BL21 | pACYC-Duet-*phoP*, pBAD-*lasR*  pCS26-*prhlR* | This study. |
| JPS2772 | *E. coli* BL21 | pACYC empty, pBAD-*lasR*  pCS26-*prhlR* | This study. |
| JPS2771 | *E. coli* BL21 | pACYC-*phoP*,pBAD empty  pCS26-*prhlR* | This study. |
| JPS2273 | *E. coli* BL21 | pACYC-*phoP* D51E, pBAD-*lasR*  pCS26-*prhlR* | This study. |
| JPS2274 | *E. coli* BL21 | pACYC-*phoP* D51A, pBAD-*lasR*  pCS26-*prhlR* | This study. |
